# Outbreak of murine infection with *Clostridium difficile* associated with the administration of a pre- and peri-natal methyl-donor diet

**DOI:** 10.1101/565762

**Authors:** Theresa Mau, Samantha S. Eckley, Ingrid L. Bergin, Katie Saund, Jason S. Villano, Kimberly C. Vendrov, Evan S. Snitkin, Vincent B. Young, Raymond Yung

## Abstract

Between October 2016 and June 2017, a C57BL/6J mouse colony that was undergoing a pre- and peri-natal methyl-donor supplementation diet intervention to study the impact of parental nutrition on offspring susceptibility to disease was found to suffer from an epizootic of unexpected deaths. Necropsy revealed the presence of severe colitis, and further investigation linked these outbreak deaths to a *Clostridium difficile* strain of ribotype 027 we term 16N203. *C. difficile* infection (CDI) is associated with antibiotic use in humans. Current murine models of CDI rely on antibiotic pretreatment to establish clinical phenotypes. In this report, the *C. difficile* outbreak occurs in F1 mice linked to alterations in the parental diet. The diagnosis of CDI in the affected mice was confirmed by cecal/colonic histopathology, presence of *C. difficile* bacteria in fecal/colonic culture, and detection of *C. difficile* toxins. F1 mice from parents fed the methyl-supplementation diet also had significantly reduced survival (*p*<0.0001) than F1 mice from parents fed the control diet. When we tested the 16N203 outbreak strain in an established mouse model of antibiotic-induced CDI, we confirmed that this strain is pathogenic. Our serendipitous observations from this spontaneous outbreak of *C. difficile* in association with a pre- and perinatal methyl-donor diet suggest the important role diet may play in host defense and CDI risk factors.

**Importance:** *Clostridium difficile* infection (CDI) has become the leading cause of infectious diarrhea in hospitals worldwide, owing its preeminence to the emergence of hyperendemic strains, such as ribotype 027 (RT027). A major CDI risk factor is antibiotic exposure which alters gut microbiota, resulting in the loss of colonization resistance. Current murine models of CDI also depend on pretreatment of antibiotics of animals to establish disease. The outbreak we report here is unique in that the CDI occurred in mice with no antibiotic exposure and is associated to a pre- and peri-natal methyl-supplementation donor diet intervention study. Our investigation subsequently reveals that the outbreak strain we term 16N203 is an RT027 strain, and this isolated strain is also pathogenic in an established murine model of CDI (with antibiotics). Our report of this spontaneous outbreak offers additional insight into the importance of environmental factors, such as diet, and CDI susceptibility.

## Introduction

*Clostridium difficile* is a spore-forming, gram positive obligate anaerobe that has become the leading cause of infectious diarrhea in hospitals worldwide. On a yearly basis, nearly half a million cases of *Clostridium difficile* infection (CDI) are reported in the United States (US), with an approximated 29,000 CDI-related deaths (1). Exposure to *C. difficile* can have varied outcomes ranging from asymptomatic intestinal colonization, to severe diarrhea, development of pseudomembranous colitis, and death (2). CDI risk is associated with disruption of the gut microbiota, for example following antibiotic administration (2–4), that leads to a loss of resistance towards *C. difficile* colonization.

Following spore germination and establishment of the vegetative form of *C. difficile*, disease results from the production of the large clostridial toxins TcdA and TcdB (5). These enterotoxins inactivate small GTPases (6–9) (e.g. Rho, Rac, Cdc42) and lead to the rearrangement of the actin cytoskeleton of intestinal epithelial cells. Consequently, mucosal epithelium damage ensues from cellular apoptosis (10) and the induction of colitis via toxin-mediated inflammation. In humans, the increasing clinical presence of *C. difficile* epidemic strains in the US and West Europe (11) is attributed to antibiotic-use and the rise of endemic strains such as NAP1/027/BI. Ribotype 027 (RT027) strains produce higher quantities of TcdA and TcdB (12) and express an additional *C. difficile* transferase (CDT) binary toxin encoded by *cdtA/cdtB* (13). The binary toxin also disrupts the cytoskeleton and increases bacterial adhesion to intestinal epithelial cells (14). The exact role of CDT in *C. difficile* virulence remains undefined, but it has been suggested that CDT plays a significant role in worsened clinical patient outcomes (15) and increases of CDI recurrence (16).

Mouse models have been developed to study the pathogenesis of CDI. These models generally require administration of antibiotics to disrupt the microbiota prior to *C. difficile* exposure. A number of antibiotic regimens have been employed to render animals susceptible to CDI (17–19). The importance of the indigenous microbiota in mediating colonization resistance against CDI is highlighted by the fact that germ free animals are inherently sensitive to colonization and disease when exposed to *C. difficile* (20).

Here, we report a spontaneous outbreak of *C. difficile* due to a strain of RT027 that occurred in a mouse colony that was associated with the administration of a specific pre- and peri-natal diet (in the original study, a methyl-supplementation donor diet (21–24) is administered to F0 mice and we study diet-induced obesity in F1 mice). Given that CDI mouse models typically rely on prior gut microbiota disruption via antibiotics to establish disease (25), this outbreak is unique in that it occurred in the absence of antibiotic use and instead, in association with an altered diet.

## Materials and Methods

### Animals

The animal care and use program at the University of Michigan is AAALAC-accredited. All procedures involving the animals and their care were approved by the University of Michigan Institutional Animal Care and Use Committee. Mice were housed in autoclaved positive-pressure individually-ventilated cages (P/NV IVC, Allentown, Allentown, NJ) with corncob bedding (The Andersons, Frontier) and provided with reverse osmosis-deionized (RO/DI) water through automated water systems (Edstrom; Waterford, WI). Animal housing rooms were maintained on a 12:12-h light:dark cycle, relative humidity of 30-70%, and temperature of 72 ± 2°F (22.2 ± 1.1°C). For specific-pathogen free (SPF) rooms, cage changing and experimental procedures were performed under a laminar flow cage change station (AniGARD^®^ VF, Baker Company, Sanford, ME) or in laminar flow benches, using a cold sterilant (Spor-Klenz^®^, Steris, St Loius, MO) for disinfecting gloved hands or transfer forceps. For biocontainment rooms (animal biosafety level 2), procedures were performed under a biosafety cabinet (SterilGARD^®^, Baker Company, Sanford, ME) using a sporicidal disinfectant cleaner (Perisept, Triple S 48027). Health surveillance program for SPF colonies included quarterly testing of dedicated soiled-bedding sentinel animals via fecal and perianal swab PCR or serology, and PCR of exhaust plenum swabs for fur mites. Health surveillance results indicated that the mice were negative for mouse rotavirus, mouse hepatitis virus, minute virus of mice, ectromelia virus, Theiler mouse encephalomyelitis virus, lymphocytic choriomeningitis virus, mouse adenovirus, mouse parvovirus, mouse polyoma virus, pneumonia virus of mice, reovirus, Sendai virus, *Mycoplasma pulmonis*, pinworms (*Syphacia* spp. and *Aspicularis* spp.) and fur mites (*Myobia musculi, Myocoptes musculinis*, and *Radfordia affinis*).

### Dietary Study

In the initial dietary study, young adult (8 wk) C57BL/6J mice were purchased from Jackson Laboratories. All custom diets “TD” are produced and distributed by Harlan-Teklad (Madison, WI). Mice were acclimated for 2 weeks before feeding either control (TD.06689) or methyl-donor supplementation (MS) diet (TD.110144). The MS diet contained 12g/kg methionine, 16.5 g/kg choline, 15 g/kg betaine, 16.5 mg/kg folic acid, 1.56 mg/kg vitamin B12, and 200 mg/kg Zn. Two weeks after starting diet, female and male mice were paired for mating. Mated F0 mice were fed the diet throughout pregnancy and lactation. F1 mice were weaned at 28 days. The F1 mice were then placed on a standard chow diet (PicoLab Laboratory Rodent Diet 5L0D, LabDiet, St. Louis, MO) or a 42% high-fat diet (HFD) (TD.88137).

### Outbreak necropsy and assessment

Dead or clinically affected animals were detected during routine daily health checks. Veterinary staff assessed for any clinical sign of gastrointestinal or systemic disease, such as diarrhea, lethargy or unkempt appearance, and euthanized animals via CO2 inhalation followed by induction of bilateral pneumothorax. Complete necropsy was performed with select tissues processed for histologic examination.

Fecal PCR of selected mice in the colony was performed by a commercial laboratory (Charles River Laboratories, Wilmington, MA) to confirm SPF status. Fecal and/or cecum and colonic samples were submitted for ELISA for *C. perfringens* type A toxin and *C. difficile* toxins A and B (Veterinary Diagnostic Laboratory, Michigan State University, Lansing, MI) or for real-time fluorogenic PCR for *C. difficile* targeting the 23S rRNA gene (Charles River Laboratories, Wilmington, MA). Cecum and colon tissues as well as tissues from other organs were fixed in 10% neutral buffered formalin for a minimum of 24 hours and then routinely processed to paraffin, sectioned, and stained with hematoxylin and eosin by the University of Michigan In Vivo Animal Core (IVAC) histology laboratory. A board-certified veterinary pathologist blinded to the diet groups evaluated the tissues descriptively in the initial outbreak and subsequently performed severity scoring of cecal and colonic tissues according to our previously published scoring system for experimentally-induced *C. difficile*-associated typhlitis and colitis (25). In brief, slides were evaluated on a 0-4 scale for the individual parameters of edema, inflammation, and epithelial damage and an overall severity score was generated by summing these parameters (scale 0-12).

To investigate the outbreak source and the extent of contamination, environmental and fecal pellet PCR was performed as described above. Swabs were taken from various areas including cardboard rolls collected for mouse enrichment and the interior of the respective collection bins and supply and exhaust plenums of four autoclaved racks. Within the ABSL2 room, door knobs, biosafety cabinet used for changing cages, clean lixits, and rack supply ducts connected to the building ventilation system were sampled. Each PCR sample tested for the affected mouse colony comprised of pooled fecal pellets (a fecal pellet collected from a representative mouse in each cage in each rack row) combined with the associated rack row’s exhaust plenum swab.

### *C. difficile* strain isolation, growth conditions, and colony identification

*C. difficile* strain 16N203 was isolated from feces frozen at −80°C after collection from a spontaneously affected animal in the dietary study mouse colony. 1.5 fecal pellets were thawed, passed into an anaerobic chamber, and diluted in 200μL sterile anaerobic PBS. 20μL was placed into 3mL of taurocholate cefoxitin cycloserine fructose broth (TCCFB) and incubated at 37°C overnight. A 10μL loop of the TCCFB enrichment culture was streaked onto taurocholate cefoxitin cycloserine fructose agar (TCCFA) to get an isolated colony of *C. difficile*. The isolated colony was used for downstream applications.

For colony identification, a 16N203 *C. difficile* colony was diluted in 15μL UltraPure Water (Invitrogen 10977-015), heated to 95°C for 20 min and then used for colony PCR to determine identity and toxin type of the isolated organism (26, 27). PCR was performed using the following primers for the *C. difficile* specific band F: 5’-TTGAGCGATTTACTTCGGTAAAGA-3’ and R: 5’-CCATCCTGTACTGGCTCACCT-3’ along with the universal 16S primers 515F: 5’-GTGCCAGCMGCCGCGGTAA-3’ and E939R:5’-CTTGTGCGGGCCCCCGTCAATTC-3’ (26). Toxin specific PCR followed previously published primers omitting the 16s rDNA primers (27). The PCRs were run in a total volume of 25uL containing GoTaq Green Master Mix (Promega M712), primers, and nuclease free water.

### *C. difficile* cytotoxin assay

Cytotoxicity assay was performed as previously described with the following modifications (25). Briefly, ATCC CCL-81 Vero cells were grown to confluence in Dulbecco modified Eagle medium (Gibco 11965) supplemented with 10% fetal bovine serum (16140) and 1% Penicillin streptomycin (Gibco 15070). Cells were plated to a density of 10^5^ cells/well. Mouse cecal content was diluted 1:10 in sterile PBS, passed through a 0.22μm filter, and serially diluted to 10^−6^. Filtered samples were tested in duplicate with a corresponding control which both antitoxin (Techlab T5000) and sample were added. A positive control of *C. difficile* TcdA (List Biologicals 152C) was used. Samples were incubated overnight at 37°C and the cytotoxic titer was determined as the reciprocal of the highest dilution that produced 80% cell rounding.

### Genome sequencing, variant identification, and comparative genomics

A colony of *C. difficile* 16N203 was cultured 18 hours anaerobically at 37°C in 13mL Brain Heart Infusion Broth (BD 211059) + 0.01% L-cysteine (Sigma C6852). The culture was spun at 4500g for 12min, washed one time with sterile PBS, spun at 4500g for 12 min and the pellet was re-suspended in 300uL of Dneasy UltraClean Microbial kit Microbead solution (Qiagen 12224-50). The extracted DNA was then prepared for sequencing on an Illumina MiSeq instrument using the NexteraXT kit and sample-specific barcoding. Library preparation and sequencing were performed at the Center for Microbial Systems at the University of Michigan. Quality of reads was assessed with Fastqc (28), and Trimmomatic (29) was used for trimming adapter sequences and low quality bases. Genome assemblies were performed using Spades (30). Variants were identified by: 1) mapping filtered reads to the assembled *C. difficile* strain 630 reference sequence (NC_009089.1) using the Burrows–Wheeler short-read aligner (BWA), 2) discarding PCR duplicates with Picard, and 3) calling variants with SAMtools and bcftools. Variants were filtered from raw results using GATK’s VariantFiltration (QUAL > 100, MQ > 50, > 10 reads supporting variant, FQ <0.025). In addition, a custom python script was used to filter out single nucleotide variants that were: 1) <5 bp in proximity to indels 2) <10 bp in proximity to another variant or 3) not present in the core genome. Maximum likelihood trees were constructed using core genome variants among the outbreak strain and a representative set of previously sequenced *C. difficile* genomes in FastTree (31). MLST predictions were made by BLASTing genome assemblies against the PubMLST database for *C. difficile* (downloaded on January 3^rd^, 2017).

### Experimental *C. difficile* infection

8-wk old C57BL/6J wild-type (WT) mice (6 males and 3 females) were obtained from an in-house breeding colony that was established originally with animals from Jackson Laboratories (Bar Harbor, ME). Mice were treated 10 days with 0.5g/L cefoperazone (MP Biomedicals 219969505) in sterile distilled water (Gibco 15230-147) as previously described (25). Briefly, animals were allowed to drink antibiotic amended water *ad libitum* and the antibiotic water was changed every other day. After 10 days, antibiotic water was switched to sterile distilled water. After 2 days on water without antibiotics, 3 male and 3 female mice (total treated n=6) were challenged via oral gavage with 500 spores of *C. difficile* strain 16N203. Spores (preparation described below) were suspended in 50μl Gibco sterile distilled water and 3 male mice (total mock n=3) were treated with 50μl of Gibco water only. Mice were monitored 16 hours post infection and feces were collected and plated to confirm spore inoculation. Mice were observed every 3 hours for 36 hours for clinical signs of disease until clinical signs appeared. Mice were euthanized by CO2 inhalation when clinical signs appeared or at study endpoint. Mice were then evaluated at 48 hours after *C. difficile* infection for weight loss, activity, posture, coat, diarrhea and nose/eyes appearance (32). Animals were euthanized and cecal contents and tissues from the animals were collected for culture and ELISA.

### *C. difficile* spore preparation

16N203 was streaked onto TCCFA and an isolated colony was incubated overnight anaerobically at 37°C in 2mL Columbia broth (BD 294420). The next day the culture was placed into 40mL of Clospore (33) and incubated anaerobically at 37°C for 8 days. The culture was spun at 3200rpm for 20min at 4°C. The pellet was washed two times with sterile water (Gibco 15230-147), once with sterile 0.05% Tween 20 (Fisher BP337), and then once with sterile water. The final pellet was resuspended in 1mL sterile water. The spore stock was stored at 4°C in sterile water. Prior to gavage, spores were heated to 65°C for 20min. Spores were plated on TCCFA to determine dose administered to animals.

## Results

### Recognition of an outbreak of severe typhlocolitis in a cohort of mice undergoing pre- and peri-natal diet manipulation

Typhlocolitis was observed in a cohort of F1 mice from a study involving parental (F0) pre- and peri-natal methyl-donor supplementation (MS) and post-weaning intake of normal diet (ND) or 42% high-fat diet (HFD) chow. The purpose of the original study was to evaluate MS diet epigenetic effects on obesity development in later-life and at various life stages of the F1 mice.

Over the course of 254 days from October 4^th^, 2016—June 15^th^, 2017, 57 out of 207 mice in the MS diet study were found dead with no premonitory signs (n=39) or were euthanized for moribund condition (n=18), including hunched posture, lethargy, dyspnea, and mucoid stool with perineal staining. Severe necrotizing typhlocolitis with pseudomembrane formation and severe submucosal edema was observed at necropsy in early cases and was suggestive of *C. difficile* colitis (25). This led to screening of affected animals for *C. difficile*, which was subsequently identified in a number of these mice, suggesting a potential outbreak. To facilitate investigation of this outbreak, we developed the following case definition: confirmed cases had 1) presentation of a compatible clinical syndrome and either, 2) histopathologic cecal/colonic lesions consistent with *C. difficile* infection or, 3) tested positive for *C. difficile* bacteria or *C. difficile* toxin. Of the 57 animals that died during the study period, we were able to complete analysis on 36 mice and determined 25 of these mice fit our case definition. We found 12 mice have causes of death unrelated to the outbreak and 20 mice had no tissues available for examination (**Table 1**).

**Table 1.** Enumeration of outbreak confirmed and excluded deaths.

|  | *Total* | *Dead* | *Examined,*<br>*Met case*<br>*definition\** | *Examined,*<br>*Did not*<br>*meet case* | *No tissue*<br>*available* |
| --- | --- | --- | --- | --- | --- |
| **All F0 mice | 24 | 1 | 0 | 1 | 0 |
| All F1 mice | 183 | 56 | 25 | 11 | 20 |
| **Control diet** (fed to F0) F1 mice | 97 | 15 | 3 | 8 | 4 |
| **MS diet** (fed to F0) F1 mice | 86 | 41 | 22 | 3 | 16 |
| **Normal chow** (fed to F1) F1 mice | 118 | 38 | 22 | 5 | 11 |
| **HFD chow** (fed to F1) F1 mice | 65 | 18 | 3 | 6 | 9 |

\*Met case definition either with histopathology or presence of *C. difficile* or *C. difficile* toxins.
\*\*F0 mice not included in survival curve analysis.

We constructed an epidemic curve to follow the course of this outbreak. The initial 14 deaths in the colony that led to the outbreak investigation were included in this epidemic curve (**Figure 1A**), but as most of these animals were found dead, or intestinal tissue was not examined, case definitions could not be assigned to these mice. Over the course of the outbreak study, there were 86 MS diet and 97 control diet F1 mice. We excluded all F0 mice and F1 mice that had no tissue available or were examined but did not meet case definition and generated a Kaplan-Meier survival curve analysis on *only the confirmed cases* of CDI shows lower survival in the MS diet mice (n=66) compared to control diet mice (n=85) (**Figure 1B**). Curve analysis indicates that on day 254, which is the day on which the last outbreak case was identified, 96.5% of the control mice survived while only 66.7% of the MS mice remained (*p* < 0.0001). A curve comparison of the confirmed CDI cases based on F1 diets—HFD (n=50) versus ND (n=101) group suggests that HFD fed mice had higher survival percentages (*p* < 0.02), (**Figure 1B**); however, there were 65 F1 mice on HFD mice while 118 F1 mice were on ND in the initial colony (**Table 1**). For comparison, we also included a Kaplan-Meier survival curve analysis on all F1 mice, incorporating them as censored (lost) for analyses (**Figure 1C**).

**FIG1.**
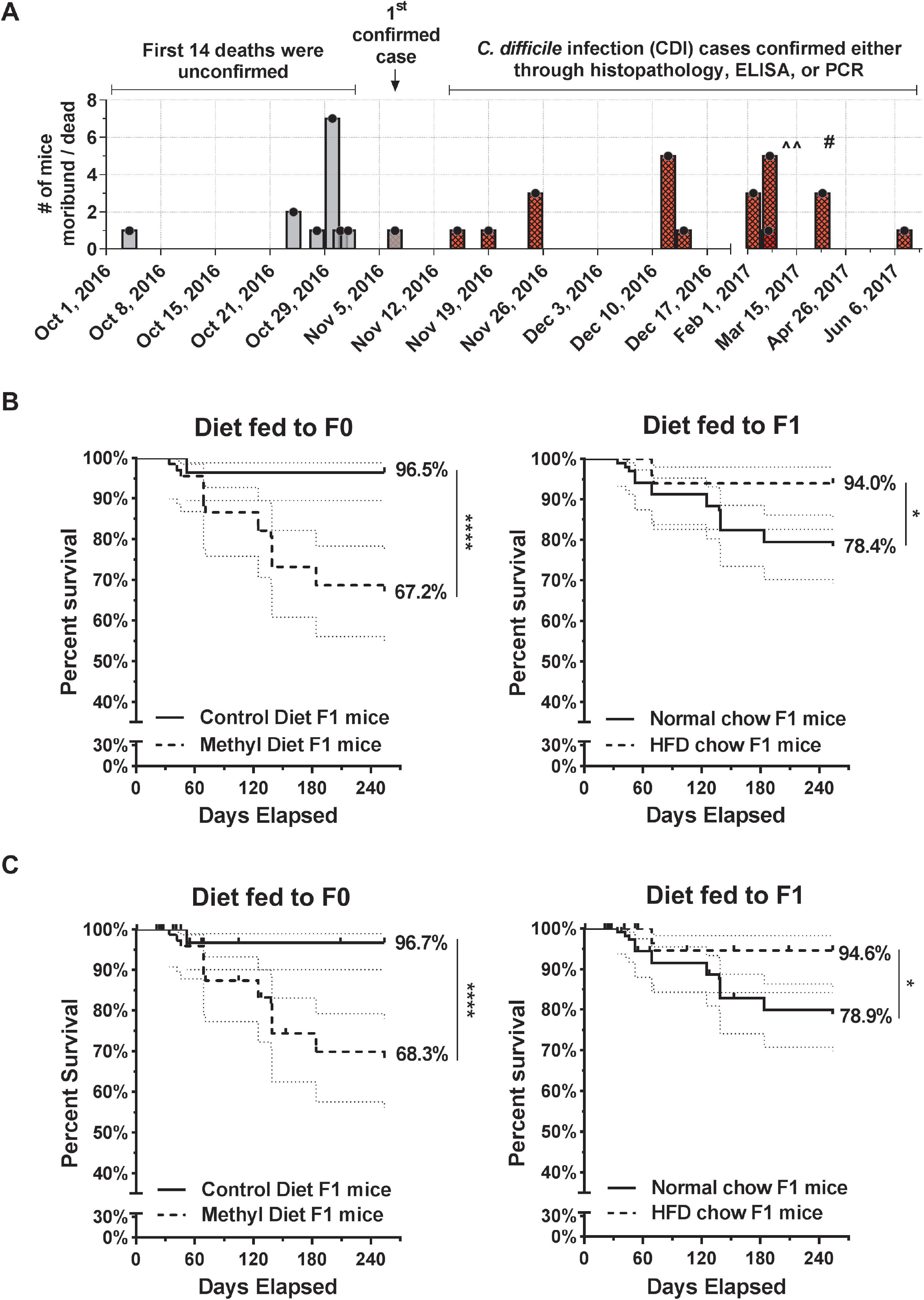
Methyl-donor diet mice had higher total mortality during outbreak. **A)** Epidemic curve depicting the first set of unconfirmed cases that triggered the outbreak investigation and the subsequent cases determined to be consistent with *C. difficile* infection.^ - *denotes environmental swab tests on 3/9/17, 3/15/17, 10/24/17 (not shown). # - denotes plenum + fecal pellet testing on 4/14/17*. **B)** Kaplan-Meier survival curves of mouse colony comparing the percent survival of the pre-/perinatal control diet (n=85, excluding 8 for not meeting case definition and 4 for unavailable tissues) versus pre-/perinatal methyl-supplementation diet (n=66, excluding 3 for not meeting case definition and 16 for unavailable tissues) F1 mice; F1 normal chow diet (n=101, excluding 5 for not meeting case definition and 11 for unavailable tissues) versus 42% high fat diet (n=50, excluding 6 for not meeting case definition and 9 for unavailable tissues) mice over the course of 8.5 months. 95% CI are graphed (light dashed lines). **C)** Kaplan-Meier survival curves including all F1 mice (n=183) in analyses but entering previously excluded mice as “0” censored. Survival curves compare Control diet (fed to F0) F1 mice (n=97) to MS diet (fed to F0) F1 mice (n=86). A second curve compares normal chow (fed to F1) F1 mice (n=118) to 42% HFD chow (fed to F1) F1 mice (n=65). * *p*<0.05, **** *p*<0.0001, Mantel-Cox log-rank test.

While attempting to control this outbreak, breeders on the MS diet and control diet were transferred to a separate room in January 2017 based on negative fecal ELISA results. The remainder of the colony was ultimately transferred to containment housing. On 4/14/17, PCR of pooled plenum and fecal pellets revealed 8/8 rows on one rack and 1/2 rows on a second rack were positive for *C. difficile*. Environmental samples on 3 separate dates (3/9/17, 3/15/17, and 10/24/17) tested negative for *C. difficile*. The timeline of the plenum swabs, fecal sampling, and environmental tests is also included on the epidemic curve (**Figure 1A**).

### Identification of *C. difficile* in animals with typhlocolitis

Gastrointestinal (cecum and colon) tissues were available for histologic evaluation in 24 of the total 57 animals found dead or euthanized during the outbreak (**Figure 2**). 17 of 24 animals were found to have necrotizing and pseudomembranous typhlocolitis consistent with CDI as seen in experimental murine models (25). Histological findings included striking submucosal edema of the cecum and colon, accompanied by marked neutrophilic inflammation and necrosis and sloughing of the superficial to mid-portions of the mucosa, often with pseudomembrane formation (**Figure 2A, 2B**). Of the 17 animals with histologically evident typhlocolitis, we were also able to collect and submit fecal and/or cecum and colonic samples from 14 animals for testing. 12 of the 14 animals tested positive for *C. difficile* TcdA and TcdB via ELISA and 2 tested positive for *C. difficile* bacteria via PCR. 14 were F1 mice from the MS diet and 3 were from the control diet groups (**Figure 2C**). The cecum summary scores (**Figure 2C**) in the MS diet (n=13) was 6.1 ± 2.6 and 9.3 ± 2.1 in the control diet (n=3). The colon summary scores (**Figure 2C**) shows that MS diet (n=14) was 4.0 ± 2.2 and control diet (n=2) was 4.5 ± 2.1. One MS-diet F1 mouse had missing colon scores and one control diet F1 mouse had missing cecum scores due to tissue autolysis. Given that significantly fewer control diet F1 mice (*p*<0.0001) were affected in the outbreak (**Figure 1B**), we collected fewer control samples and thus cannot draw conclusions on the effect of the F0 diet on disease severity between the two groups of F1 mice.

**FIG2.**
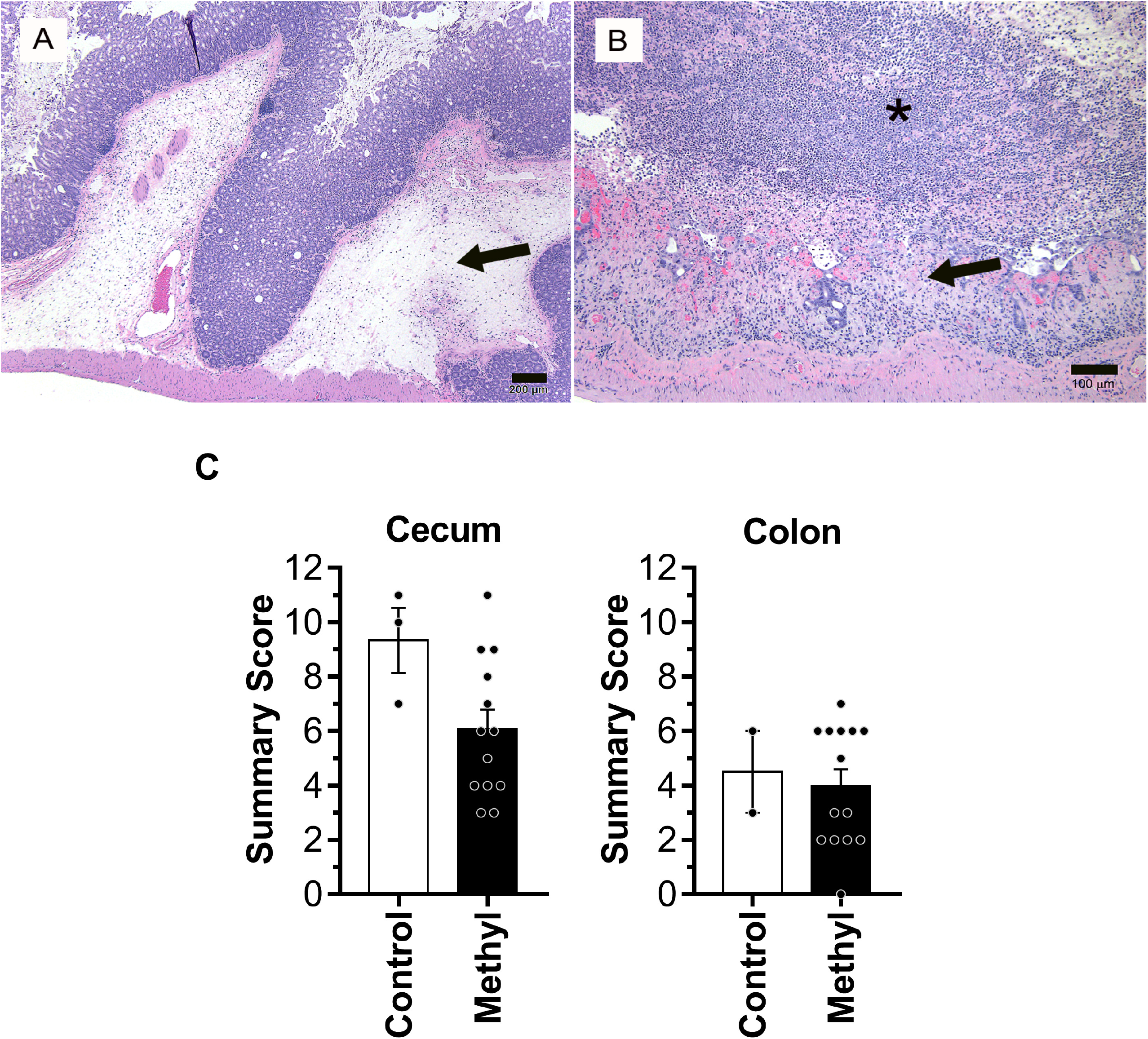
Necropsies and histopathology indicate typhlocolitis as primary diagnosis of affected animals. **A)** Representative histology of mouse cecum in a *C. difficile* culture-positive case exhibiting severe submucosal edema and inflammation (arrow). H&E stain, Bar=200 μm. **B)** Representative histology of an area of cecum in the same case displaying mucosal necrosis (arrow) and extensive pseudomembrane formation (*), consisting of neutrophils and sloughed enterocytes in a fibrinous matrix. H&E stain, Bar=100 μm. **C)** Histological severity scores for the 17 mice showing typhlocolitis, out of 24 clinically affected mice for which tissues were available. Severity was scored by a board-certified veterinary pathologist using an established murine *C. difficile* scoring system. Each category—edema, inflammation, or epithelial damage was individually scored 0-4 and a summary score was generated (range 0-12). Of the 17 mice, 1 mouse had no cecum score and 1 mouse had no colon score due to tissue autolysis of those samples. As such, for cecum scores, control diet F1 mice n=3 and methyl-supplementation diet (MS) F1 mice n= 13. For colon scores, control diet n=2 and MS diet n=14.

Overall, we observed that the histology score was higher in cecum than colon in most animals, consistent with experimental murine *C. difficile* models (25). Of the remaining 7 out of 24 histologically evaluated animals that did not have evidence of typhlocolitis, alternate causes of death/morbidity were identified, including tumors (n=2), bacterial enteritis (n=2), or undetermined by histology (n=3). Finally, there were 12 out of 36 analyzed animals that had no gastrointestinal tissue available to be histologically evaluated, and 8 of these animals tested positive for *C. difficile* bacteria (n=2) and *C. difficile* toxins A and B (n=6) which then allows us to include these 8 as confirmed cases of *C. difficile-related* deaths. The 4 animals that tested negative for *C. difficile* toxin were found to have an alternative cause of death (endometritis n=1) or undetermined (n=3), and naturally, these were not included in the confirmed cases of *C. difficile*.

Isolation of *C. difficile* was performed by the use of selective culture. A *C. difficile* strain (designated 16N203 for the animal from which it was initially isolated) was obtained in pure culture. A PCR (26) confirmed the *C. difficile* bacteria was present in fecal sample (**Supp. Figure 1A**), and this strain was also positive for the toxin genes *tcdA/tcdB* and *cdtA/cdtB* by PCR (27) (**Supp. Figure 1B**). Whole genome sequencing of strain 16N203 was undertaken, and a whole genome phylogeny was constructed (**Figure 3A**), including a representative set of previously sequenced *C. difficile* isolates. These isolates contain publicly available clinical genomes (34–36), clinical isolates collected at the University of Michigan, and two mouse strains (16N203 and LEM1). LEM1 is an indigenous murine spore-forming *C. difficile* strain identified and isolated from mice acquired from common mouse vendors, Jackson Laboratories and Charles River Laboratories (37). LEM1 appears to not be highly virulent and can protect against the closely related but more virulent strain VPI10463, at least in mice with C57BL/6J or BALB/c background (37). In our data, the murine strain LEM1 clusters with Clade 1, which contains reference strain CD630, while the outbreak strain fell within the diversity of Clade 2, which contains RT027 isolates (**Figure 3A**). When a phylogenetic tree was constructed with only RT027 isolates, the outbreak 16N203 strain clustered with isolates derived from human patients with clinical CDI (38) (**Figure 3B**).

**FIG3.**
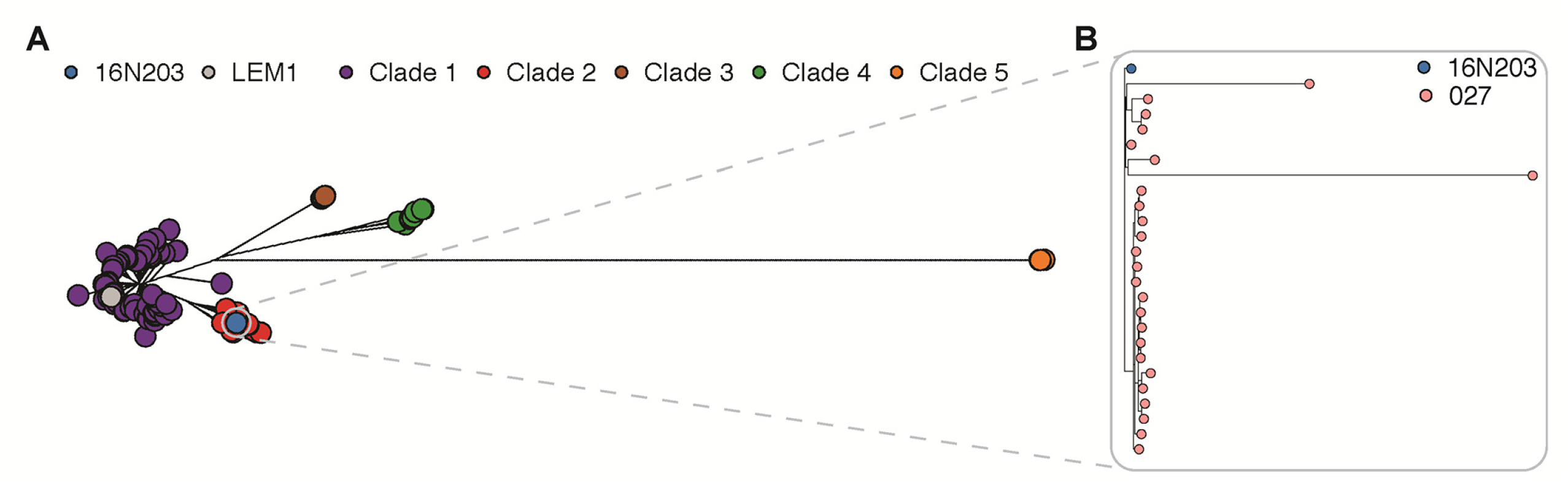
Isolation and genomic sequencing confirms NAP1/027/BI *Clostridium difficile* strain. **A)** Phylogenetic relationship between the mouse outbreak strain and representative *C. difficile* isolates. Isolates include publicly available clinical genomes, clinical isolates collected at the University of Michigan, and two mouse strains (16N203 and LEM1). A maximum likelihood tree was generated from the recombination filtered, polymorphic core genome positions in all isolates (N = 438). Clade 1 (purple) includes reference strain CD630 and mouse strain LEM1 (gray); clade 2 (red) includes epidemic RT 027. Clades 1-5 are as previously defined (36, 54, 55). **B)** Tree in (A) was subset to include only 027 isolates. The mouse outbreak “16N203” strain (blue) clusters with the RT 027 isolates (pink), a RT in Clade 2 (red).

### *C. difficile* strain 16N203 is fully virulent in a mouse model of antibiotic-induced CDI

We had previously demonstrated that *C. difficile* strains have variable virulence in an established mouse model where CDI susceptibility is conveyed by treatment with the antibiotic cefoperazone (25). To determine whether the pathogenicity of the outbreak strain 16N203 was unique to the dietary model, we assessed whether the outbreak strain 16N203 was virulent in this established model of CDI (**Figure 4A**). C57BL/6J mice on a standard diet and without previous dietary manipulations were treated with cefoperazone in drinking water for 10 days and then switched back to plain water. Two days after stopping antibiotics, mice were challenged with either 500 spores of the 16N203 outbreak strain or vehicle via oral gavage. Significant weight loss (*p*<0.05) was observed in mice challenged with 16N203 spores during the 48-hour study time course while mock-treated mice maintained weight (**Figure 4B**). Mice challenged with *C. difficile* 16N203 had higher levels of cecal colonization (*p*<0.03) with detectable toxin in intestinal contents (*p*<0.05) and developed clinical signs compatible with severe CDI (**Figure 4C**).

**FIG4.**
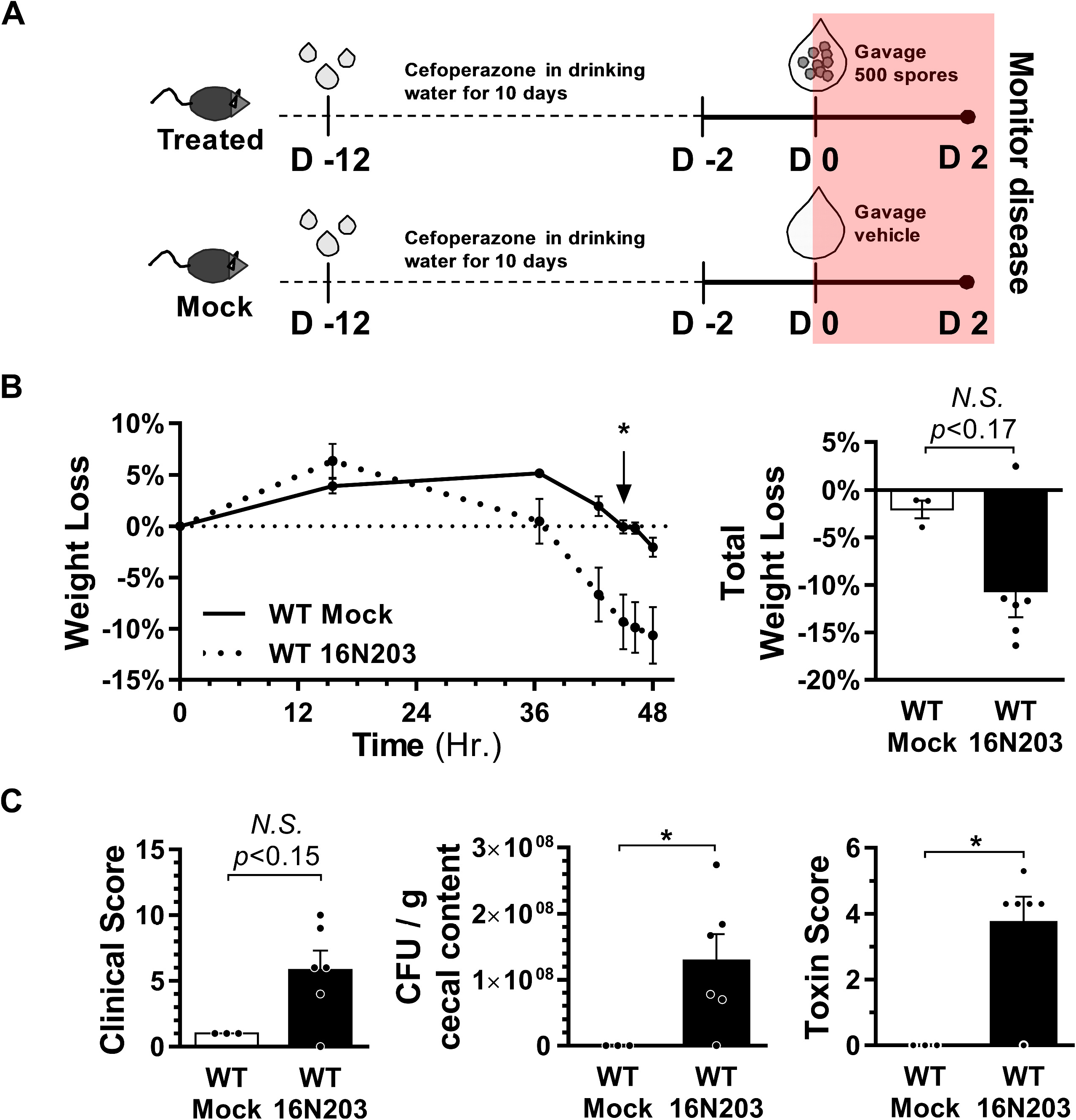
*C. difficile* ‘outbreak’ strain induces CDI in a standard mouse model with antibiotics. **A)** CDI mouse model with cefoperazone in drinking water used to assess virulence of the outbreak 16N203 strain. Wild-type (WT) mice were administered cefoperazone in drinking water for 10 days on day −12 (D −12) and then orally gavaged either 16N203 spores (n=6) or the vehicle (distilled water) for mock (n=3) controls on day 0 (D0). Mice were monitored for disease between D0 and D2. **B)** Body weight loss over the course of 48 hours; total percentage of body weight loss shown in column graph. **C)** Clinical score, colonization, and toxin scores at 48 hours. For weight loss, clinical, colonization, and toxin scores, Mann-Whitney test was used for statistical analyses, * *p*<0.05.

## Discussion

In this report, we describe an outbreak of *C. difficile* infection (CDI) and colitis in laboratory mice associated with diet manipulation. This outbreak is interesting because spontaneous, symptomatic CDI is unusual in laboratory mice, and symptomatic colitis due to experimental CDI generally requires pretreatment with antibiotics (37, 39). Subclinical colonization of mice has been reported previously. In one study it was noted that mice could be colonized with *C. difficile* with or without prior antibiotic treatment but this colonization was at a very low level and not associated with any clinical disease (19). Administration of the antibiotic clindamycin was associated with increases in the levels of colonization but again without development of clinical colitis. It has been also reported that wild rodents can harbor toxigenic *C. difficile* strains, again without any overt disease (40). One group reported that laboratory mice obtained from multiple sources appear to harbor LEM1, an indigenous *C. difficile* strain (37). Colonization with LEM1 in the same study did not result in disease and in fact was associated with protection from disease following challenge with another strain of *C. difficile* that generally produces severe colitis in experimental models of CDI. In contrast, while LEM1 clusters with Clade 1 (containing reference strain CD630 (RT 12)), the outbreak strain in our study clustered in Clade 2 with patient-derived RT 027 isolates and was associated with clinical disease in both spontaneously infected and experimentally infected mice. The source of this strain remains unclear, and our attempt at revealing a source with environmental sampling did not yield conclusive results. Ribotyping and genetic analyses suggest a potential source was through fomites or human carriers, but the source may have been gone by the time of environmental investigation.

The mechanism underlying the unexpected increased susceptibility of these mice is also not entirely clear. Small animal models to recapitulate CDI pathogenesis generally rely on antibiotic pretreatment prior to experimental challenge with *C. difficile*, even with virulent strains (39). As noted, the current outbreak is unusual in that there were no antibiotics administered to the animals and instead susceptibility to colonization and colitis was associated with dietary (peri- and pre-natal) manipulation. Our current understanding of the pathogenesis of CDI attributes loss of colonization resistance to altered structure and function of the indigenous intestinal microbiota (41). However, given the unexpected nature of the outbreak, we were not positioned to determine if there were changes in the microbiota associated to the increased susceptibility we observed in MS F1 mice. Antibiotics, which can cause widespread changes in the indigenous microbiota, are associated with the development of clinical CDI in humans, but other factors that alter microbial populations may also convey susceptibility.

In our study, increased *C. difficile*-associated deaths were seen in the offspring of mice receiving the methyl-supplementation (MS) diet. Diet may be a factor that can alter the community structure and function of the intestinal microbiota, and diet-related susceptibility to CDI has been evaluated in some studies (42, 43). Patients receiving enteral tube feeding had increased risk of developing CDI (44). Tube feeds given in the form of an elemental diet (i.e. a diet that is entirely absorbed within the small bowel) are thought to deprive the colonic bacteria of nutrition in the form of fiber and resistant starch, increasing the risk of CDI (45). Microbiota accessible starches may protect against CDI as in murine studies where feeding such carbohydrates can suppress experimental CDI (46). This protection was associated with an increase in short chain fatty acids which are the metabolic products of microbial fermentation of non-digestible carbohydrates. Another study showed that a low protein diet (which had reciprocal increases in carbohydrate composition) could be protective in a mouse model of CDI (47). In addition to macronutrients, it was recently demonstrated that zinc deficiency could alter the microbiota and increase susceptibility to experimental CDI (48).

It should be noted that, in contrast to our study, the diet-related susceptibility or amelioration of experimental CDI discussed above occurred in models where the microbiota was still altered via antibiotic administration. An earlier study in hamsters fed an atherogenic diet in the absence of antibiotic administration, unexpected diarrhea and death due to colitis was observed in animals starting 45 days after diet manipulation (49). Similar to our study, toxigenic *C. difficile* was isolated from affected animals, although typing of the strain was not performed. It is interesting to note that in these hamsters, the atherogenic diet increased susceptibility to CDI, but in our study, F1 mice that were on a 42% high fat diet (HFD) exhibited increased survival—though it should be noted that this observation was limited by the fact that there were significantly more normal diet mice than HFD mice in the mouse colony at the time.

While hamsters are exquisitely susceptible to *C. difficile*, symptomatic murine *C. difficile* colitis outbreaks in the absence of antibiotic administration are very rare. There is precedence for disease outbreaks in mice associated with *C. difficile*, but the affected mice have had significant immune alterations that increased their susceptibility (50, 51). One such outbreak occurred in a murine experimental autoimmune encephalomyelitis model generated via the administration of pertussis toxin and a myelin oligodendrocyte glycoprotein fragment (51). These authors speculated that the stress of this experimental manipulation or the development of the autoimmune disease altered the microbiota in a manner that leads to susceptibility to CDI, but they did not profile the microbiota in these animals. Although we were unable to characterize the microbiota in our study due to the unexpected nature of the outbreak, a future avenue to explore would be diet-associated microbial community alterations that may convey susceptibility.

Specific mechanisms by which our methyl-donor diet alters susceptibility to CDI are unclear. A similar methyl-supplementation maternal diet was shown to increase colitis risk in a chemically-induced inflammatory bowel disease (IBD) using dextran sulfate sodium (DSS) (52, 53). F1 mice from methyl-fed dams had worsened colitis and significantly higher mortality than control F1 mice (52). Colonic mucosal bacterial diversity analyses indicated striking composition variation, including significantly higher prevalence of *Clostridia* in methyl diet F1 mice. In addition, there was an increase in *Firmicutes (Lachnospira, Oscillospira, Ruminococcus* and *Catonella*) and a decrease in *Parabacteroides* and *Eubacterium* in methyl F1 mice compared to controls. This suggests that dysbiosis between colitogenic (*Firmicutes*) and anti-inflammatory bacteria (*Parabacteroides*) following prenatal methyl supplementation may have led to the colitis prone phenotype (52). A fecal microbiome transfer (FMT) via cage swapping (between methyl F1 mice and germ free mice) was sufficient to worsen DSS-induced colitis in germ free mice, indicating that the maternal methyl-donor diet alone may enhance a colitis-prone microbiota profile. While this study cannot be directly compared to *C. difficile* colitis, it strongly suggests that a prenatal methyl-donor diet is sufficient in generating a pro-colitic F1 microbiota. In the context of CDI, it remains to be determined whether susceptibility is due strictly to altered community composition, creating a niche for *C. difficile* colonization, or whether microbial changes directly promote colitis-enhancing inflammatory responses. While unexpected, the current spontaneous outbreak of CDI in the absence of antibiotics in mice with parents fed an altered pre- and peri-natal diet may ultimately provide additional insight into the interplay of diet, host responses, and the microbiota that mediates CDI and colitis susceptibility.

## Acknowledgements

The authors would like to thank Aline Penkevich for helping with the cytotoxin assays of *C. difficile*, the staff of the In Vivo Animal Core for histology work, and the Rodent Health Surveillance Team at the Unit for Laboratory Animal Medicine for environmental investigation of the outbreak.

---

**Supplementary FIG 1. PCR identification of *Clostridium difficile* and its associated toxins (*TcdA, TcdB*, and the binary *cdtA/cdtB* toxin) A)** PCR (26) confirming *C. difficile* specific identification. Lane 1 = 100bp ladder, Lane 2 = 16N203-1, Lane 3 = 16N203-2, Lane 4 = 16N203-3, Lane 5 = 16N203-4, Lane 6 = *C. difficile* 630, Lane 7 = water. **B)** PCR (27) confirming presence of CDI toxins: Toxin A (*TcdA*), Toxin B (*TcdB*), and binary toxin (*cdtA/cdtB*). Lane 1 = 100bp ladder, Lane 2 = 16N203-1, Lane 3 = 16N203-1, Lane 4 = 16N203-2, Lane 5 = 16N203-3, Lane 6 = 16N203-4, Lane 7 = *C. difficile* R20291, Lane 8 = *C. difficile* 630.

